# Reducing Pollen Dispersal using Forest Windbreaks

**DOI:** 10.1101/204263

**Authors:** Thomas Meyer, Vernie Sagun, Carol Auer

## Abstract

The adoption of genetically engineered (GE) crops has created a demand for practical methods to mitigate pollen dispersal and gene flow. The goal of this project was to measure the ability of a narrow forest windbreak to reduce downwind pollen fluxes from switchgrass (*Panicum virgatum* L.), a North American grass and model biofuels feedstock. Switchgrass fields were established in two identical plots where one had a forest windbreak and the other was in an open (control) site. Switchgrass reproduction, pollen dispersal, wind speed, and wind direction were measured over two years. Daily release of switchgrass pollen peaked at 11:00-13:30 during a flowering period that lasted about 44 days. The best estimate for switchgrass pollen source strength (PSS) was 141 × 10^9^ pollen/season/hectare for fields planted at commercial densities. The forest windbreak consistently decreased downwind switchgrass pollen concentrations by 333-20,000 fold compared to the control plot which had a 58-77 fold decrease due to downwind distance alone. These results suggest that forest windbreaks could be used as a barrier to reduce pollen dispersal and gene flow from switchgrass and other crops.

**Research Highlight:** A narrow forest windbreak greatly decreased downwind pollen concentrations from a switchgrass field suggesting that trees can reduce crop gene flow, enhance coexistence between farming systems, and provide ecosystem services.

## 1. Introduction

Plant gene flow, the movement of genes from one plant population to another, is a natural part of plant reproduction and speciation [Ellstrand, 2014]. However, pollen-mediated gene flow from genetically engineered (GE) crops has created huge challenges for farms with organic certification, producers of identity-preserved crops, seed companies that must maintain genetic purity, companies producing non-GE food products, and government agencies regulating biocontainment of experimental field trials [USDA AC21, 2012, Wilkinson and Tepfer, 2009, Andow and Zwahlen, 2006, Auer, 2008]. Furthermore, the dispersal of some transgenic pollen might negatively impact native plant communities, non-target organisms, or the broader environment over time. Unfortunately, there are few practical methods to reduce pollen dispersal and gene flow from agricultural fields. This project examined the ability of a narrow forest windbreak to reduce downwind pollen concentrations from fields of switchgass (*Panicum virgatum* L.), a lignocellulosic biofuel crop. The long-term goal was to develop tree windbreaks as a practical method of reducing pollen-mediated gene flow while enhancing ecosystem services in the landscape.

In theory, gene flow from wind-blown pollen can be reduced by geographic features [Graves et al, 2014], appropriate isolation distances [Slatkin, 1993], specific agricultural practices (e.g. barriers, trap crops, or buffer rows) [McNaughton, 1988], or genetic engineering techniques [Ellstrand and Hoffman, 1990]. At present, only isolation distances and trap crops are used to reduce transgene movement. Pollen dispersal by wind is affected by various factors including wind speed and direction [Clark et al, 2005, Ecker et al, 2013, Hoyle and Cresswewll, 2007, Jia et al, 2007, Pfender et al, 2007, Wang and Yang, 2010, Wang et al, 2004]. A Lagrangian model of switchgrass pollen dispersal under typical summer wind conditions in the Northeastern US showed that viable pollen could be transported 3.5 km – 6 km from the source field [Ecker et al, 2013]. Thus, reducing wind speeds across agricultural fields (pollen sources) could play a role in reducing crop-to-crop, crop-to-wild, or crop-to-weed gene flow. Another important factor in gene flow is pollen source strength (PSS), defined in this study as the quantity of pollen that can be dispersed per unit crop area [Muilenberg, 1995]. This study is the first to predict PSS for commercial switchgrass fields.

Windbreaks reduce wind speed, and they have been used for various applications including the protection of crops and livestock [Brandle et al, 2004]. The micrometeorological changes produced by windbreaks and forest edges have been well studied [Dupont et al, 2007, Eimern et al, 1964, Heisler et al, 1988, McNaughton, 1989, Patton et al, 1998]. Windbreaks produce turbulence in the wind field that dissipates the flow’s kinetic energy, which suppresses downwind wind speeds within a distance of 10 – 20 times of the height of the windbreak [McNaughton, 1989, Patton et al, 1998]. The effectiveness of a windbreak in creating a protected downwind zone is determined by factors including its height and porosity [Cleugh, 1998], upwind land cover, and topography.

The potential for windbreaks to reduce pollen dispersal has received little attention. Arritt et al. [2007] found that a vegetative border of a tall annual grass (sorghum-sudangrass hybrids) reduced pollen transport from a maize field. Ushiyama et al. [2010] evaluated fences as windbreaks in maize fields, but found little effect on pollen dispersal. Although research has shown that pollen concentration decreases with distance from the source [Ecker et al, 2013, Chamecki and Meneveau, 2011, Okubo and Levin, 1989], there are no studies on the effect of typical farm windbreaks (one or more rows of cultivated trees or shrubs) or narrow forest stands on pollen dispersal. In this paper, a barrier made from a narrow forest stand is called a *forest windbreak*.

Understanding and managing pollen-mediated gene flow in switchgrass is important because it is a model crop for large-scale biofuels plantations and a GE crop with novel traits approved by the US government [Ecker et al, 2015, Ledford, 2013]. Switchgrass is a native grass across the Central and Eastern regions of North America with minimal domestication from its wild ancestors. Its reproductive biology includes obligate out-crossing, self-incompatibility, numerous panicles, and a long period of asynchronous pollen release [McNaughton, 1988, Casler, 2012]. Various ecological risks from GE switchgrass have been proposed including the emergence of switchgrass as a serious weed, the reduction or extinction of native switchgrass populations, the loss of natural genetic biodiversity and genetic resources, and negative impacts on non-target organisms [Barney and DiTomaso, 2008, Kausch et al, 2010, Kwit and Stewart, 2012]. Additional concern exists because switchgrass is already a successful competitor outside of cultivation [Ecker et al, 2015, Kausch et al, 2010]. A study in the Northeastern US analyzed the genetics of switchgrass collected from road verges and the coastal zone where local populations are lowland tetraploids [Ecker et al, 2015]. Results showed that 67% of the roadside plants were upland octoploid plants that were probably introduced through human activity. Thus, mechanisms to reduce gene flow from biofuels plantations could help protect local plant communities [Barney and DiTomaso, 2008, Kausch et al, 2010, Kwit and Stewart, 2012].

The purpose of this study was to: 1) characterize switchgrass flowering, pollen release (anthesis), pollen dispersal, and pollen source strength; 2) measure the effect of a forest windbreak on wind speed and direction; and 3) determine if a forest windbreak could reduce downwind switchgrass pollen dispersal beyond what would occur due to distance alone.

## 2. Materials and Methods

### 2.1 Field Plots in the Study

The project was conducted at two 40 m × 40 m field plots in Ecoregion Level IV number 59 (Northeastern Coastal Zone [Griffith, 2010]) (Fig. 1) planted with switchgrass in 2012 with observations made at plant maturity in 2013 and 2014. One field plot at 41.792N, 72.223W was set in a second-growth forest (hereafter called “Forest”). The second field plot was at 41.781N, 72.212W in agricultural land without windbreaks (hereafter called “Control”). Soils were classified as sandy loams with low-to-medium organic-matter content with pH 5.8 – 6.

**Figure 1.**
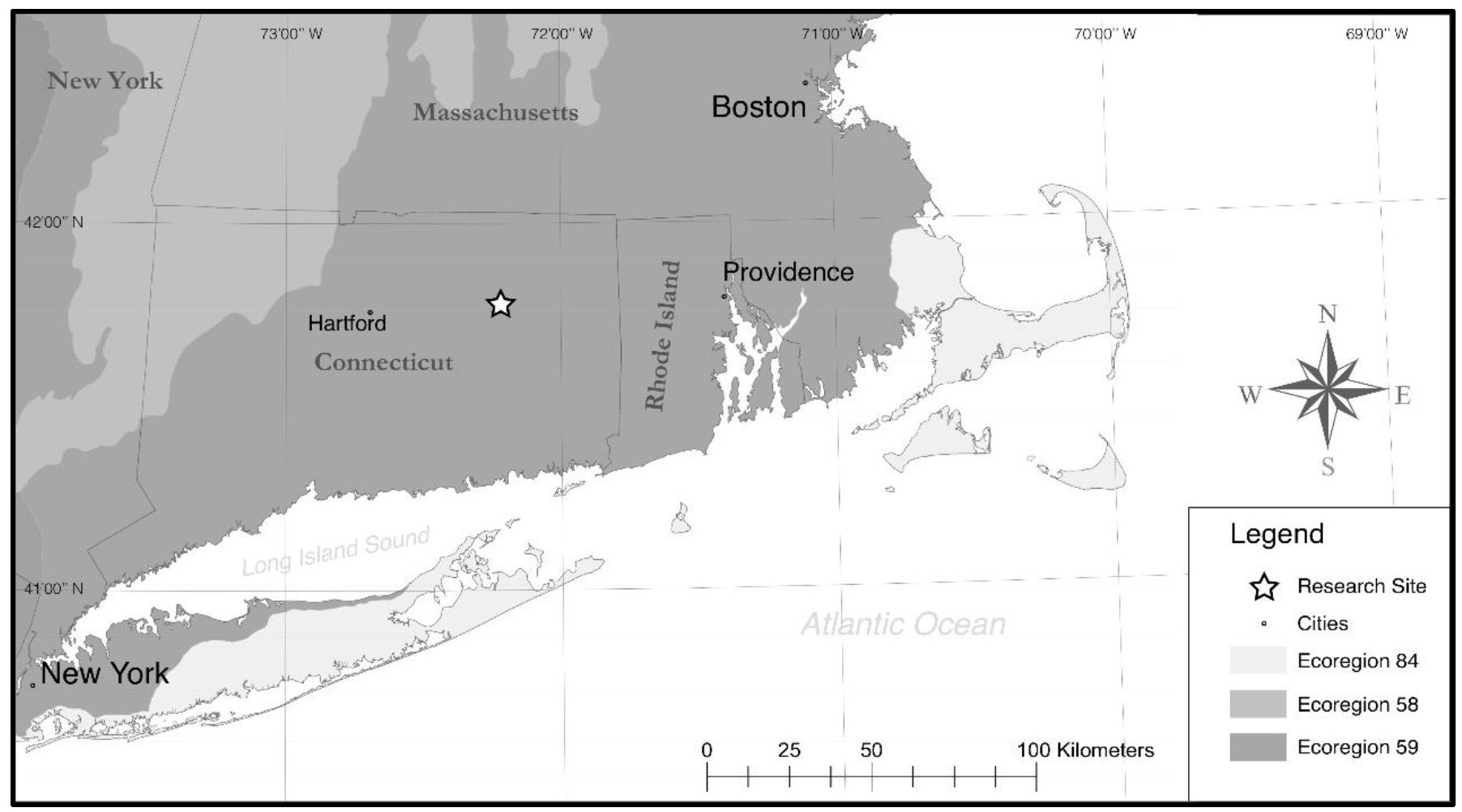
Location of field study (star) in the context of Level IV Ecoregions in the Northeastern United States. Locations are shown for four major cities (Boston, Hartford, New York City, Providence) and borders for four states (Connecticut, Rhode Island, Massachusetts, New York).

A botanical survey was conducted to characterize the second-growth forest windbreak. There were 197 trees with >4 cm diameter breast height, and the species included *Fraxinus americana* (50.7%), *Acer rubrum* (22%), *Carya ovata* (22%), *Malus* sp. (2%), *Ostrya virginiana* (1%), *Quercus velutina* (1%) and *Sassafras albidum* (0.5%). The forest understory contained a small number of shrub species including *Ilex verticillata*, *Lindera benzoin*, *Rosa multiflora* and *Berberis thunbergii*. Vines in the forest canopy included *Celastrus orbiculatus*, *Parthenocissus quinquefolia*, *Toxicodendron radicans*, and *Vitis* species.

The forest windbreak was mapped using a Leica Scan Station C10 terrestrial laser scanner (Leica Geosystems, San Francisco, California, USA). This instrument is a robotic total station that scans horizontally and vertically, collecting a *xyz* point every few centimeters including returns from the highest leaves so long as there is a direct line-of-sight to the instrument. From these data we determined mean tree height to be 16.8 m.

### 2.2 Establishment of Switchgrass Fields

Two switchgrass cultivars were planted in alternating rows: ‘Blackwell’ (upland octoploid, Natural Resource Conservation Service, US Department of Agriculture, Salina, Kansas, USA) and ‘Cave-in-Rock’ (upland octoploid, Natural Resource Conservation Service, US Department of Agriculture, Elsberry, Missouri, USA). In 2012, switchgrass seedlings were grown for 8 weeks in Fafard^®^ Nursery Potting Mix (Conrad Fafard Inc., Agawam, MA, USA), fertilized with 3 g of Nutricote^®^ 18-6-8 (type 180, Arysta LifeScience, Chuo-ku, Tokyo, Japan) and grown in a University of Connecticut greenhouse (mean temperature 26°C, range 19-38°C). Seedlings were planted in mid-June in rows 1 m apart with 0.5 m between individuals. Each row had 80 individuals for a total of 3200 plants per plot. Plant survival during the first winter (2012-2013) was approximately 98% for both fields and dead plants were replaced in June, 2013. Weeds were controlled by hand within rows and trimmed between rows. Both fields were fertilized in mid-April, 2013 with 50 kg/ha of organic Milorganite^®^ All Season Fertilizer (5-2-0) (Milorganite, WI, USA). Switchgrass plants were cut down after frost in mid-November in all years.

### 2.3 Switchgrass Development

To understand switchgrass reproduction, 20 plants were randomly selected for each cultivar and plot location (40 plants/field, 80 plants total). The timing of flower development and anthesis was determined based on weekly surveys in 2013 beginning prior to the emergence of flower panicles and continuing until all plants had finished flowering. One panicle from each plant was tagged and observed weekly during flowering to count the number of florets with exerted purple stigmas and yellow-orange anthers; these florets were designated as in the process of anthesis (releasing pollen). All 80 individuals were harvested, dried and weighed for above-ground biomass in October, 2013. Two panicles were collected from each individual and the number of florets was counted. The number of reproductive stems (with panicles) and vegetative shoots (without panicles) were counted. Analysis of Variance was used to test differences in above-ground biomass, number of reproductive shoots and number of florets/panicle. Tukey’s HSD was used to test for significant differences between cultivars and plots [R Core Team, 2013].

### 2.5 Pollen Capture

Switchgrass pollen were sampled in the air using rotorod boxes constructed by the authors. A rotorod box houses a DC motor attached to a rotating-arm impactor that has two narrow plastic impaction surfaces oriented vertically (rods) that spin in a circle [Muilenberg, 2003]. The plastic rods (2.98 mm width × 2.54 cm length on leading side) sampled a volume of air ranging from 0.70 m^3^ – 0.76 m^3^ in 30 minutes depending on the angular velocity of each rotorod box (velocities varied from 2900 – 3170 rpm). The leading side of the rods was given a thin coating of silicone grease, and rods were replaced every 30 minutes. After exposure, the rods were transported to the lab where the number of pollen were counted under an Olympus SZH10 microscope (Olympus, PA, USA) along a 1-cm length at the center of each rod. The rotorods were digitally photographed and the images stitched together using the software ImageJ 1.47v [Rasband, 2014] and counted using SemAfore 5.21 (JEOL (Skandinaviska) AB, Sollentuna, Sweden). Pollen counts were converted into number of pollen/m^3^/30 minutes with an adjustment for the speed of each individual rotorod box where the m^3^ refers to the volume of air sampled. Eight case study days with wind and pollen data were analyzed with three days from the Control site (Aug. 21, 2013 and Aug. 19 and 29, 2014) and five from the Forest site (Aug 31, 2013 and Aug. 14, 20, 25, 26, 2014).

Estimates of pollen source strength (PSS) were based on the number of pollen/anther, anthers/floret, florets/panicle, reproductive stems/plant, and the field density (number of individuals/area). To determine the number of pollen per anther, one switchgrass floret from a ‘Cave-in-Rock’ plant was collected and placed in Clarke's fixative solution (25% acetic acid to 75% ethanol). The mature flower bud was dissected and each anther was placed in 15 μL Calberla’s staining solution and 15 μL glycerol and dissected to release pollen. The pollen suspension was then placed in a Fuchs-Rosenthal haemocytometer and photographed in an Olympus BH2-RFCA microscope (Olympus, PA, USA). Pollen counts were made twice for each anther using the program SemAfore 5.21 (JEOL (Skandinaviska) AB, Sollentuna, Sweden).

Pollen production (*P*) for an individual plant was calculated with the formula

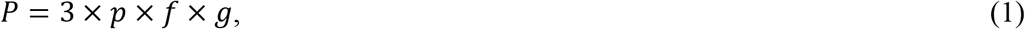

where there were three anthers per florets, *p* is the mean number of reproductive stems with panicles per plant, *f* is the mean number of florets per panicle, and *g* is the mean number of pollen per anther. The standard deviation of *P* was calculated using propagation of variance [Ghilani, 2010] as

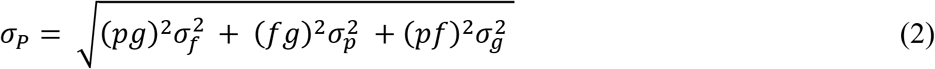

The maximum number of airborne pollen throughout the flowering period was estimated by multiplying the value *P* by the number of plants in the experimental plot or the number of plants/area (density) in a commercial field.

### 2.4 Wind Dynamics

Sonic anemometers (Ultrasonic 81000, RM Young, Michigan, USA) were deployed as shown in Figure 2 to characterize the wind fields. Three 3-m towers were erected at the Forest plot (denoted by first letter F) and the Control plot (denoted by first letter C). Each tower supported a sonic anemometer at the top of a 3-m metal pole with a rotorod box mounted at 2-m height. At each plot, one tower was erected in the middle of the switchgrass field (denoted by second letter S), one was 99.8 m from the middle of the switchgrass field (denoted by second letter D_1_), and one was 133.7 m from the middle of the switchgrass field (denoted by second letter D_2_). Thus, the six towers were named FS, FD_1_, FD_2_, CS, CD_1_, and CD_2_. The distances for D_1_ and D_2_ were chosen according to the mean forest canopy height of the Forest windbreak. Downwind distances were D_1_ = 1 × mean tree canopy height (16.76 m) and D_2_ = 3 × height (50.3 m). Thus, D_1_ = 16.8 m and D_2_ = 50.3 m from the downwind edge of the forest windbreak. To measure pollen dispersal within and above the forest windbreak, a 23.8 m walkup tower (denoted by W) was erected inside the forest 19.5 meters from the middle of the switchgrass field and in line with the 3-m towers (Figs. 2, 3). The top of the tower extended above the tree canopy by about 2 meters. Sonic anemometers and rotorod boxes were placed on the walkup tower at 1.8 meters (W_1.8_) and the top of the tower (W_23.8_). W_23.8_ was directly above W_1.8_, so they appear as W in Figure 2. A third sonic was mounted at 12.8 meters on the walkup tower (W_12.8_), but pollen was not collected at this height.

**Figure 2.**
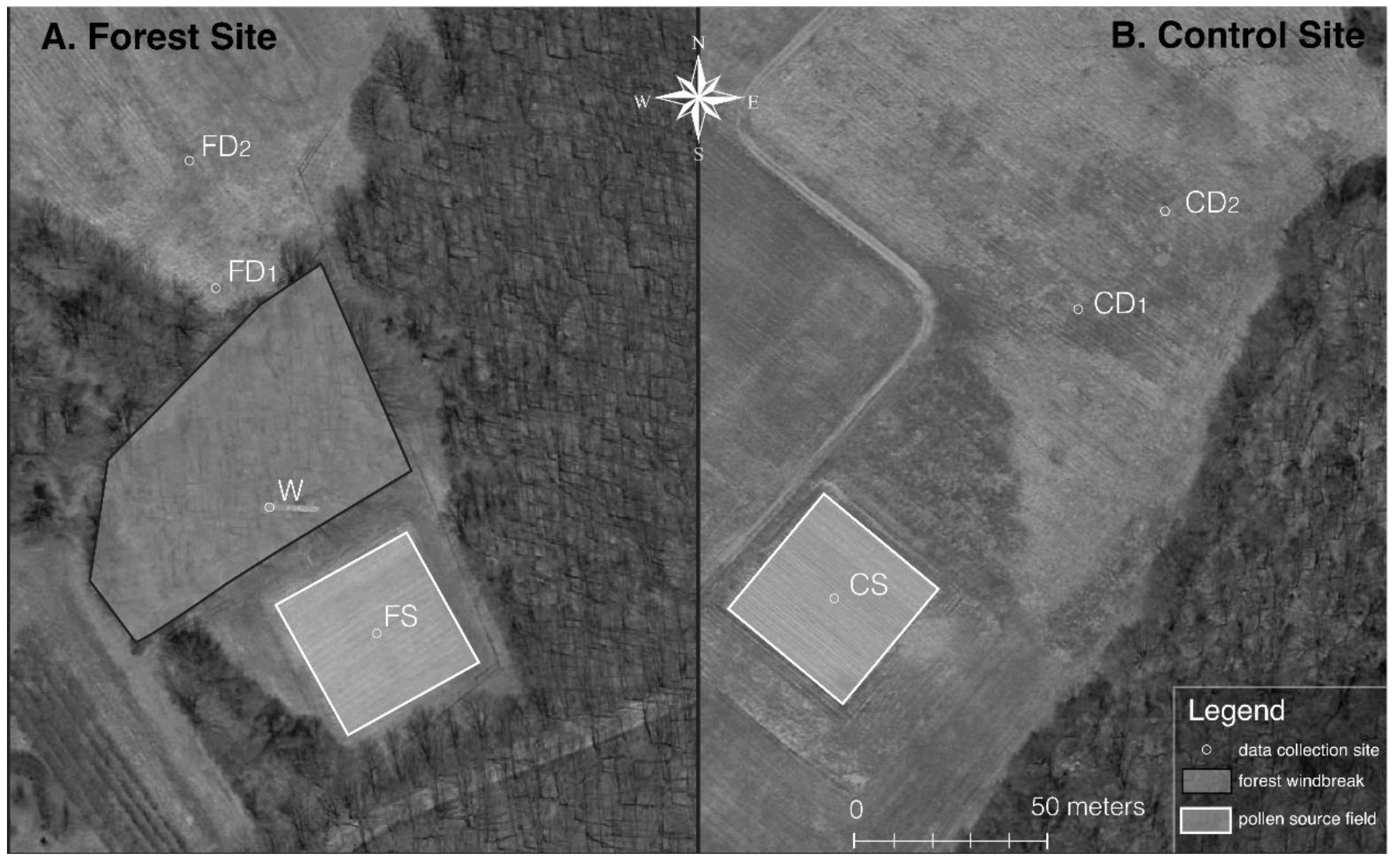
Aerial photographs of switchgrass field plots. The Forest plot (left) shows a square representing the 40 m × 40 m Forest switchgrass field (FS), a polygon representing the forest windbreak, the location of the Walkup tower in the forest (W), and two downwind data collection towers (FD_1_, FD_2_). The Control plot (right) shows a square representing the 40 m × 40 m Control Source field (CS) and two downwind data collection towers (CD_1_, CD_2_).

**Figure 3.**
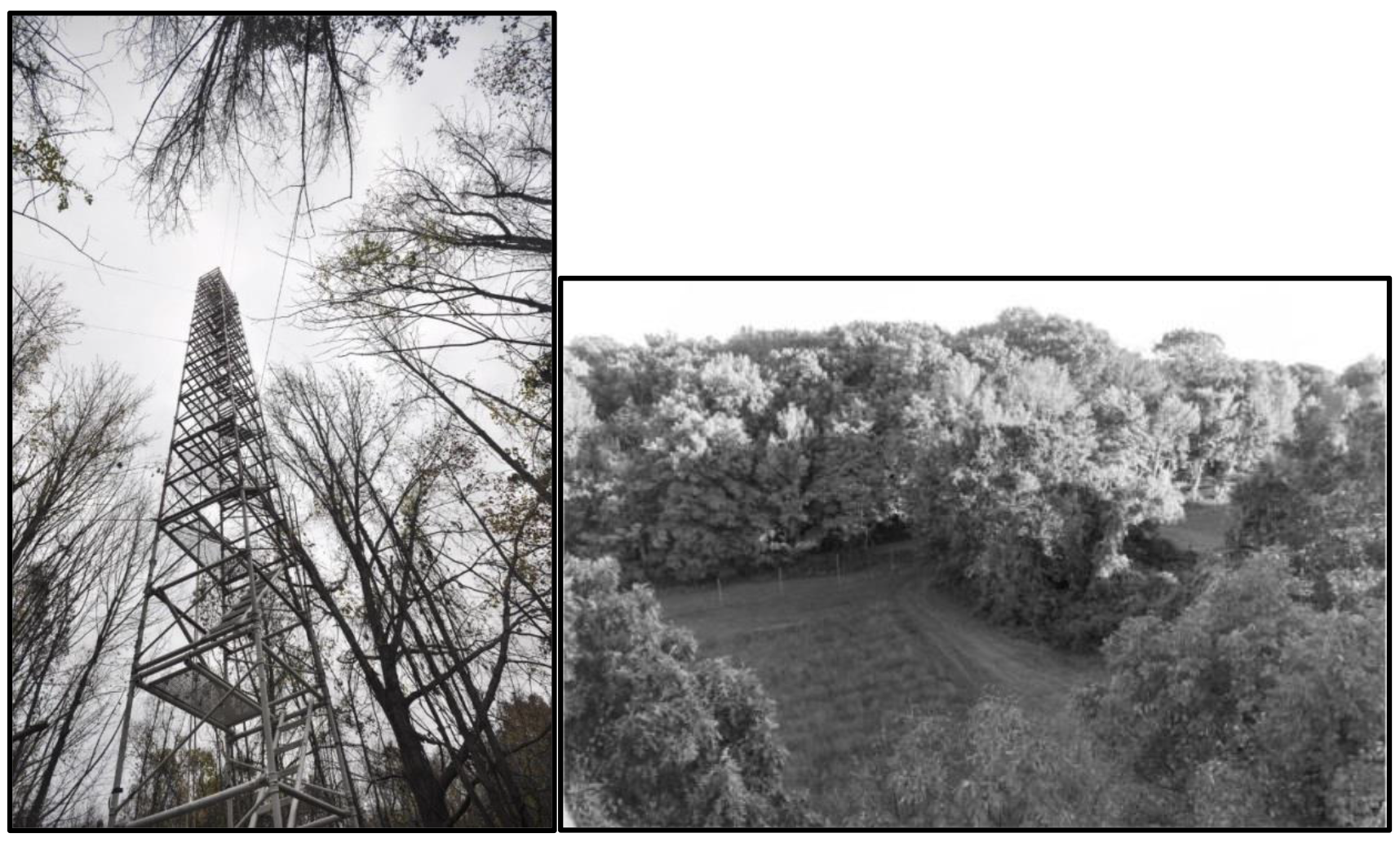
Photographs of the walkup research tower (Left) extending through the forest windbreak canopy and the switchgrass Forest Source field (Right) as seen from the walkup tower.

The sonic anemometers measured three-dimensional wind-velocity fields (m/s) and temperature (C) at 10 Hz, and were oriented with their fiducial axis aligned to the north. The two horizontal components were denoted as +u for wind blowing to the north, +*v* for wind blowing to the east, and +*w* for the vertical component blowing upwards (Fig. 4). Data were collected on netbook computers (Acer Aspire One D270, Acer America, CA, USA). The netbooks ran the Ubuntu ^TM^ 12.04 LTS Linux ^TM^ operating system and sonic data were collected with PuTTY ©, a free and open source serial console and file transfer application (http://www.chiark.greenend.org.uk/~sgtatham/putty/).

**Figure 4.**
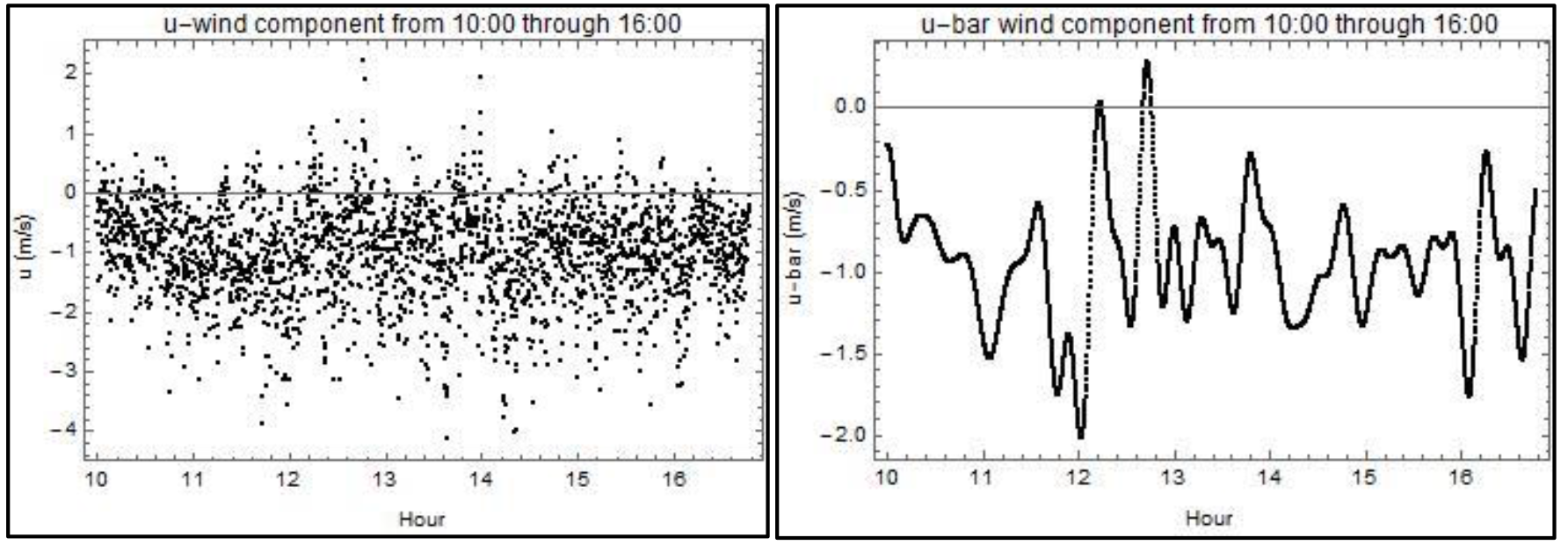
Sample of wind data collected by a sonic anemometer showing northwards wind velocity component *u* (m/s) in the control field on August 21, 2014. Left panel: Observed wind *u* component. Right panel: Mean speed 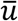. (c) Turbulence 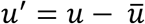. Data were subsampled by 1 in 100 to reduce clutter in the image.

## 3. Results

### 3.1 Switchgrass Anthesis

Switchgrass panicles were first observed emerging from stems on July 2, 2013 and July 11, 2014. Panicle emergence in ‘Blackwell’ (B) was slightly delayed relative to ‘Cave-in-Rock’ (CIR) in both plots. Observation of 80 randomly selected plants in 2013 showed that the pattern of anthesis was similar among cultivars and plots and occurred over approximately 44 days (Fig. 5). The number of florets in anthesis/panicle was not statistically different among cultivars and plots except on Aug. 15 and Sept. 4, 2013, but no cultivar or plot had consistently higher values on those dates. With few exceptions, there were no significant differences among cultivars and plots for above-ground biomass (477.2 grams dry weight ± 284.2 SD), number of florets/panicle (705.9 ± 284.2 SD), and number of flowering stem/individual plant (75.8 ± 33.8 SD). However, the biomass for CIR plants in the Control plot (770 g ± 285.5g) was higher than for B plants in the Forest 14 plot (270.5 g ± 157 g) and Control CIR had a slightly higher number of flowering stems than Forest CIR, Forest B or Forest CIR.

**Figure 5.**
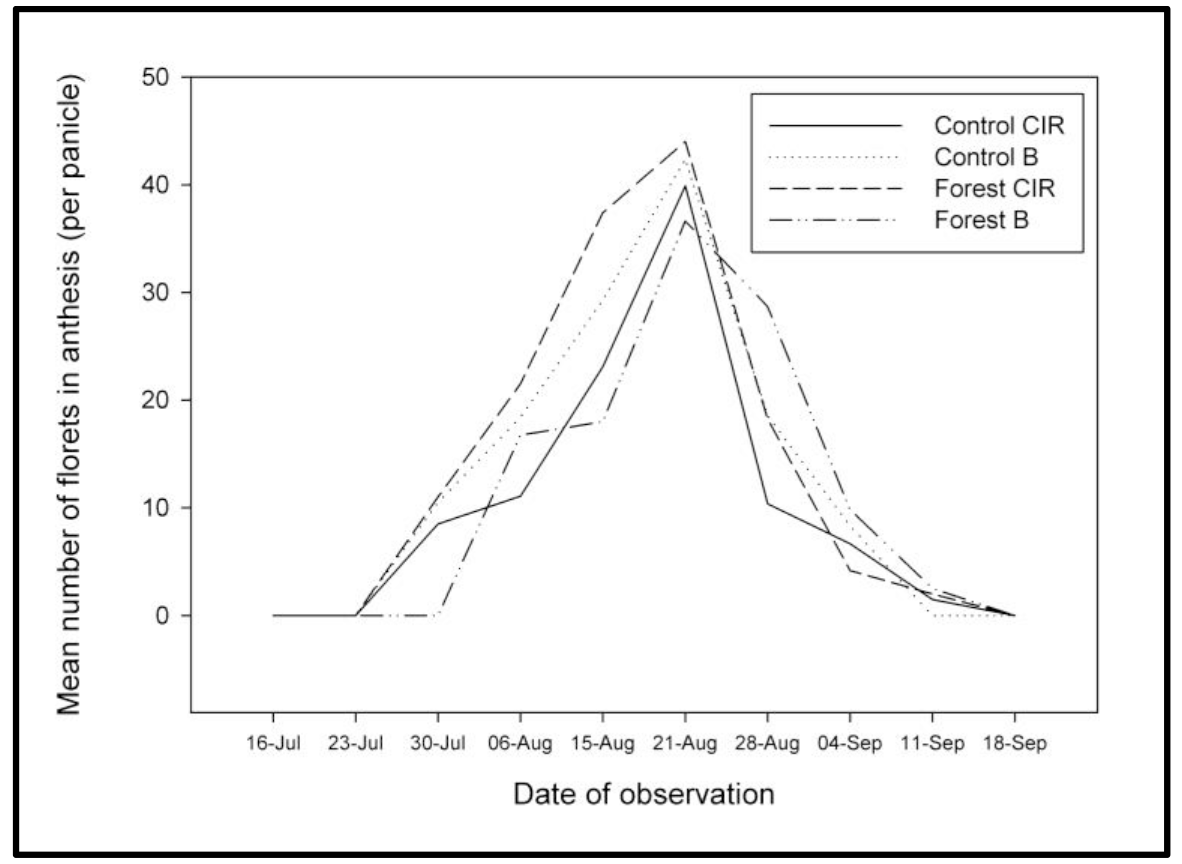
Mean number of switchgrass florets in anthesis per panicle during the flowering period (2013). Cultivars ‘Cave-in-rock’ (CIR) and ‘Blackwell’ (B) in Forest and Control plots. Twenty panicles were observed for each plot and cultivar combination (*n* = 20).

The pattern of switchgrass pollen release during the day was determined by collecting pollen on rotorods in the center of the switchgrass fields above the panicles on eight different days (Fig. 6). Pollen were first observed on rotorods at 11:00 with a broad peak in pollen concentrations at 11:30-13:30 followed by a decrease to lower levels by 15:00.

**Figure 6.**
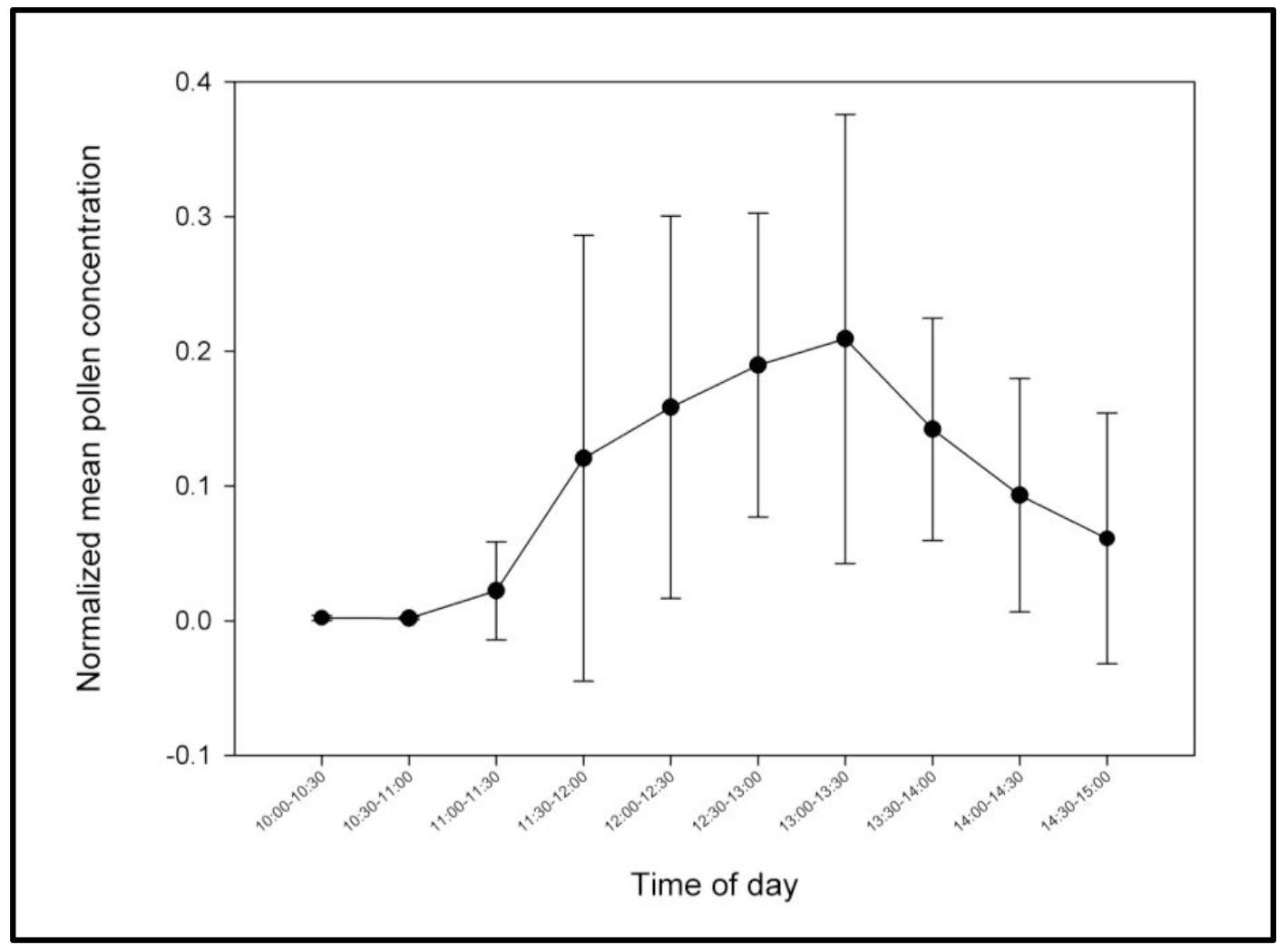
Pattern of switchgrass pollen collection during the day. Normalized mean percent values were based on the number of pollen captured in each 30 min period (pollen/m^3^/30 minutes) divided by the total number of pollen captured in that day by rotorod boxes at the center of the source fields. The normalized percent values and standard error are shown for 30 min intervals from 10:00 – 15:00 (n = 8 case study days). For example, about 20% of the pollen collected for the day was captured from 13:00 – 13:30.

### 3.2 Pollen Source Strength

The first step in estimating pollen source strength (PSS) for the switchgrass fields was to determine the mean pollen production/switchgrass plant over the growing season. This was estimated using values for the mean number of florets per panicle *f* = 706 ± 284), the mean number of panicles per plant (*p* = 76 ± 34), and the mean number of pollen per anther (g = 5500 ± 839; min = 4504, max = 6771). Using Eq.s 1 and 2, the estimated maximum number of pollen emitted per plant per growing season was 8.85 × 10^8^ ± 1.83 × 10^8^. Using this information, three values for PSS were developed: 1) PSS for 3200 plants in the experimental field plots, 2) PSS for switchgrass fields planted at commercial densities, and 3) an adjusted PSS based on pollen capture data. With 3200 plants per plot, we estimate that our field plots produced 2.83 × 10^12^ ± 1.4 × 10^10^ pollen per season per plot. Assuming commercial planting density is one plant per square foot [USDA NRCS, 2009] a commercial biofuels plantation would be approximately four times denser than our plots and a hectare is 6.25 times larger, so the estimate for total maximum source strength was 7.08 × 10^13^ ± 5.18 × 10^10^ pollen per season per hectare at commercial densities. However, the airborne PSS under natural conditions would generally be less than this because not all the pollen in the anthers is released, some pollen falls directly to the ground, some florets or anthers are damaged, and pollen number could vary due to genetic or environmental factors [Cruden, 2000]. Therefore, these values are an upper bound based on the cultivars and environment in this study.

Dividing the maximum source strength estimate for our plot by 6 weeks/season and 7 days/week gives 6.75 × 10^10^ ± 1.60 × 10^9^ pollen per day for the plot. The 40 m × 40 m plot in square centimeters is 1.6 × 10^7^ cm^2^ and a rotorod box occupies 42.25 cm^2^, so the entire field would hold a theoretical maximum of 378,698 rotorod boxes. Dividing 7.29 × 10^10^ pollen per day for the plot by 378,698 rotorod boxes suggests that each rotorod box would collect 178,119 pollen per day. In fact, the mean pollen number collected in the source field was 16,881 pollen in a day (min=5650, max=29083). Because pollen collection occurred between 10:00-15:00, it is possible that some pollen was released outside this time frame and other factors could reduce pollen cloud density (see above). Nonetheless, the maximum estimate is 11 times larger than the observed mean. Taking this into account, a more realistic peak release estimate from a commercial field is 6.75 × 10^9^ pollen/day/hectare when planted at commercial densities.

Taking 6.75 × 10^9^ pollen/day/hectare as the maximum release in a single day, we estimated the pollen/day/hectare on days other than the peak release. Table 1 gives the percent-of-maximum observed on the other observation days. The maximum occurred on 08/21 as indicated by observed percent of maximum equal to 100%. Pollen release on unobserved days was estimated by linear interpolation (Fig. 7). Summing the observed and interpolated values^1^ produces our estimated pollen source strength for the whole growing season: 141 × 10^9^ pollen/season/hectare. To our knowledge, this is the first estimate of switchgrass PSS in the literature.

**Table 1.**
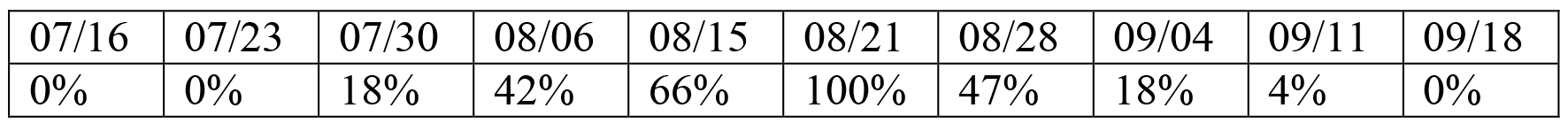
Observation dates and observed percent-of-maximum pollen released per day.

**Figure 7.**
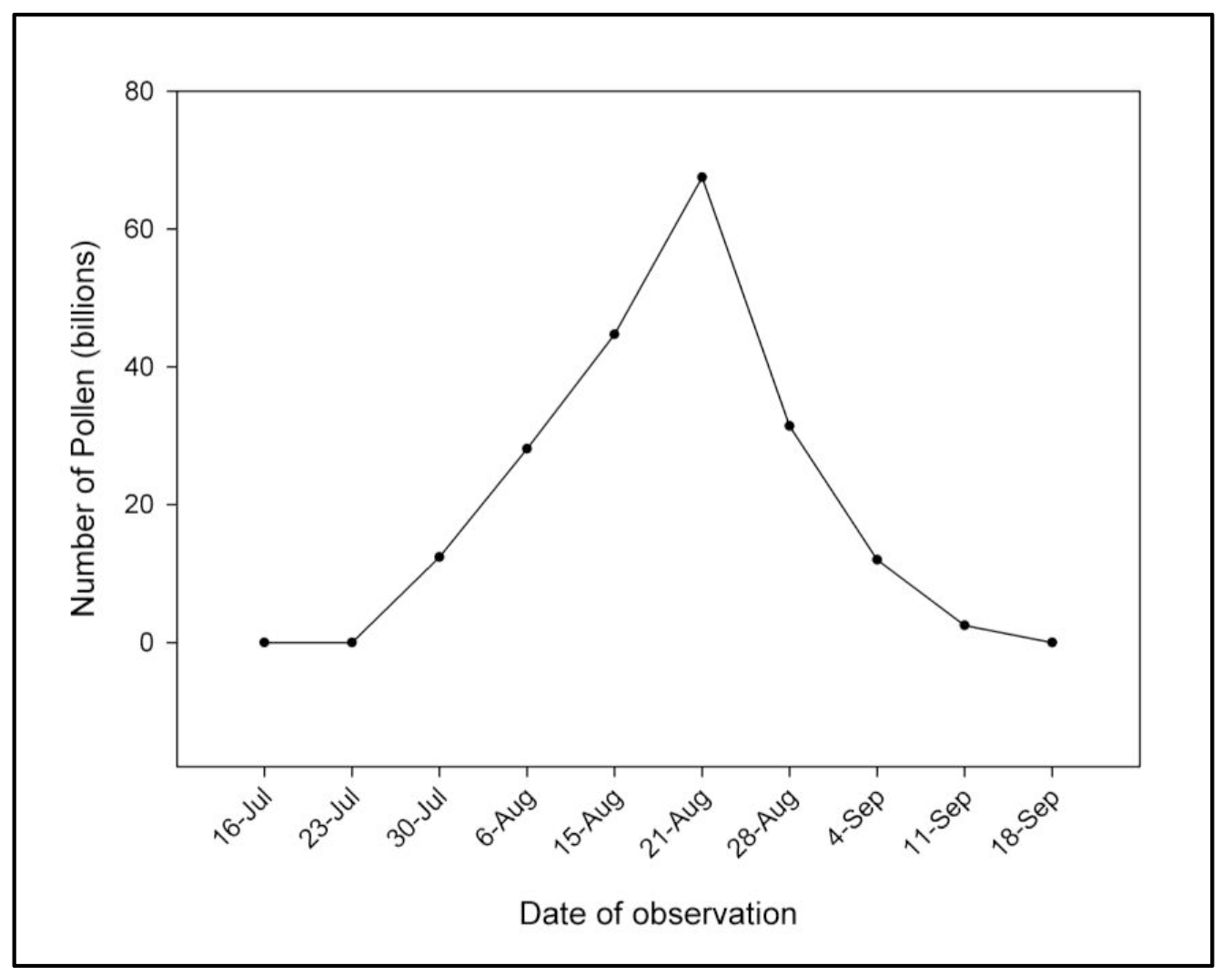
Estimation of the number of switchgrass pollen released over the growing season based on the observed number of florets in anthesis/panicle and an estimate of total pollen source strength for the switchgrass field (3200 plants).

### 3.3 Effect of the Forest Windbreak on Pollen Dispersal

Data from eight case-study days (2013-2014) are presented to understand the effects of the forest windbreak on pollen dispersal (Table 2). Most of the case-study days occurred when the principle wind direction was from the pollen source fields towards the towers with rotorod boxes and sonic anemometers (called “downwind”). One case study (August 14, 2014) at the Forest plot was conducted with the opposite trend in wind direction. Table 2 summarizes the observed pollen concentrations at each rotorod box (observed pollen numbers for the day together with those values normalized with the center of the field), wind speed, and wind direction. Pollen concentrations were always highest in the center of the source field compared to all other collection points. At both 19 plots, pollen concentrations at Downwind 2 (D2) were lower than at Downwind 1 (D1) with the exception of August 31, 2014 when the values were nearly equal (14 pollen and 18 pollen).

**Table 2.** Switchgrass pollen concentrations and wind data for eight case study days. Pollen concentrations (pollen/m^3^/5 hr from 10:00-15:00) are shown for each rotorod location with both observed values and normalized values. Wind data is shown as speed (m/s) and direction (degrees from North azimuth). Wind speeds for the Forest plot are shown at the center of source field and (/) top of walkup tower. Wind speeds for the Control plot are shown for the center of the source field.

| Plot Name | Case Study Date | Pollen Concentration (pollen/ m<sup>3</sup> / 5 hr) |  |  |  |  | Mean wind speed | Minimum wind speed | Maximum wind speed | Mean Wind direction |
| --- | --- | --- | --- | --- | --- | --- | --- | --- | --- | --- |
|  |  | Center Source Field | Top tower | Bottom tower | Downwind 1 | Downwind 2 |  |  |  |  |
| Forest plot | August 14 2014 | 5650 | 42 | 44 | 6 | 3 | 0.88/2.04 | 0/0 | 5.35/9.33 | -17 |
| Forest plot | August 20 2014 | 8135 | 426 | 282 | 23 | 13 | 0.63/1.06 | 0/0 | 2.62/4.09 | -4 |
| Forest plot | August 25 2014 | 19632 | 34 | 105 | 1 | 0 | 0.71/1.76 | 0/0 | 4.20/6.16 | 2 |
| Forest plot | August 26 2014 | 23642 | 991 | 210 | 24 | 10 | 0.66/1.01 | 0/0 | 3.15/5.53 | -27 |
| Forest plot | August 31 2013 | 9216 | 165 | 26 | 14 | 18 | 0.62/2.20 | 0/0 | 3.60/7.25 | -23 |
| Control plot | August 19 2014 | 29083 |  |  | 405 | 141 | 1.1 | 0 | 5.51 | -6 |
| Control plot | August 29 2014 | 15141 |  |  | 254 | 24 | 1.17 | 0 | 5.45 | 12 |
| Control plot | August 21 2013 | 20370 |  |  | 261 | 104 | 1.4 | 0 | 5.46 | -29 |
| Plot Name | Case Study Date | Normalized Pollen Concentration |  |  |  |  | Mean wind speed | Minimum wind speed | Maximum wind speed | Mean Wind direction |
|  |  | Center Source Field | Top tower | Bottom tower | Downwind 1 | Downwind 2 |  |  |  |  |
| Forest plot | August 14 2014 | 100.0 | 0.743 | 0.779 | 0.106 | 0.053 | 0.88/2.04 | 0/0 | 5.35/9.33 | -17 |
| Forest plot | August 20 2014 | 100.0 | 5.237 | 3.467 | 0.283 | 0.160 | 0.63/1.06 | 0/0 | 2.62/4.09 | -4 |
| Forest plot | August 25 2014 | 100.0 | 0.173 | 0.535 | 0.005 | 0.000 | 0.71/1.76 | 0/0 | 4.20/6.16 | 2 |
| Forest plot | August 26 2014 | 100.0 | 4.193 | 0.887 | 0.102 | 0.042 | 0.66/1.01 | 0/0 | 3.15/5.53 | -27 |
| Forest plot | August 31 2013 | 100.0 | 1.795 | 0.286 | 0.150 | 0.194 | 0.62/2.20 | 0/0 | 3.60/7.25 | -23 |
| Control plot | August 19 2014 | 100.0 |  |  | 1.393 | 0.485 | 1.1 | 0 | 5.51 | -6 |
| Control plot | August 29 2014 | 100.0 |  |  | 1.678 | 0.159 | 1.17 | 0 | 5.45 | 12 |
| Control plot | August 21 2013 | 100.0 |  |  | 1.282 | 0.520 | 1.4 | 0 | 5.46 | -29 |

Pollen concentrations at the Control plot consistently decreased with distance from the center of the switchgrass field: there was a 58-77 fold decrease in pollen concentrations from the center of the source field to CD_1_ (99.8 m from field center) and 200-1000 fold decrease at CD_2_ (133.7 m from field center). These reductions are explained by distance alone because the control site had no barriers between the source field and the downwind sampling towers. The Forest windbreak plot consistently decreased the downwind pollen concentrations far more than what was observed at the Control plot. Pollen concentrations at FD_1_ (beyond the Forest windbreak) decreased by 333-20,000 fold, and pollen concentrations at FD_2_ decreased 500-2500 fold. No switchgrass pollen was detected at FD_2_ on one case study day (Aug 25, 2014). Pollen concentrations were also measured at two positions on the forest Walkup tower. Pollen concentrations above the forest tree canopy (W_23.8_) were 19 - 500 fold less than the center of the switchgrass field, and at the bottom of the Walkup tower (W_1.8_) they were 29 – 333 fold less than the center of the switchgrass field. However, the ratio of pollen at the top (W_23.8_) and bottom (W_1.8_) rotorod boxes varied between days. Overall, it is clear that downwind pollen concentrations (CD_1_, CD_2_) were always higher at the Control plot.

Results for two days (Figure 8) were chosen for comparison because both fields had comparable pollen source strength: 23,642 pollen/m^3^/5 hr at the Forest plot and 29,083 pollen/m^3^/5 hr at the Control plot. The pollen collector beyond the forest windbreak (FD_2_) had no detectable pollen (0%) compared to 0.5% at the same distance in the Control site (CD_2_); similar differences were observed at FD_1_ and CD_1_.

**Figure 8.**
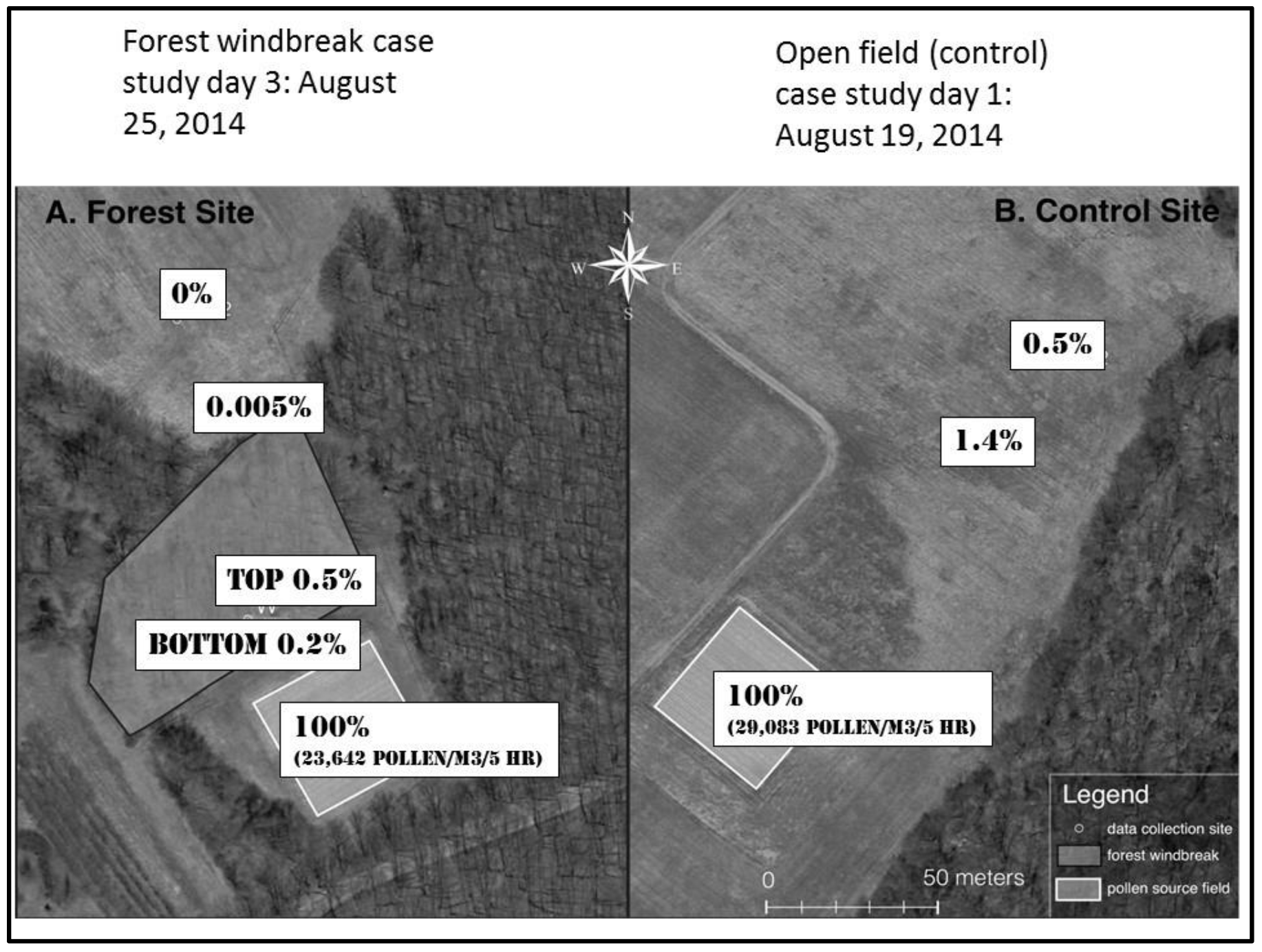
Two aerial photographs showing reduction in pollen concentrations downwind from the source fields. The left photo shows normalized pollen concentrations at the Forest plot. The pollen collector at the top of the walk-up tower (W_23.8_) above the forest canopy is labeled ‘TOP’ and the collector at the bottom of the tower (W_1.8_) is labeled ‘BOTTOM’. The right photo shows pollen concentrations at the Control plot.

### 3.4 Micrometeorology

Wind direction is an important factor in pollen dispersion [Clark et al, 2005, Hoyle and Cresswell, 2007, Okubo and Levin, 1989, Augspurger and Franson, 1987]. Wind roses were computed from the anemometer observations for every 30 minute pollen sampling period, and some “typical” exemplars are shown. The sector gray tones reflect mean wind velocity towards that direction for that half hour, with darker colors indicative of higher wind speeds relative to 3 m/s. The length of a sector is in proportion to the number of samples blowing that direction, so the wind rose gives the same information as a frequency plot (histogram). Figure 9 shows wind roses for 10:00-10:30 at the Control plot on August 29, 2014. The wind fields at the three sonic anemometers are very similar in direction and speed. They are well organized and consistent. The downwind sensors recorded higher average wind speeds than the sensor in the middle of the field, which is reasonable because the vegetation around the field sensor slows the wind somewhat.

**Figure 9.**
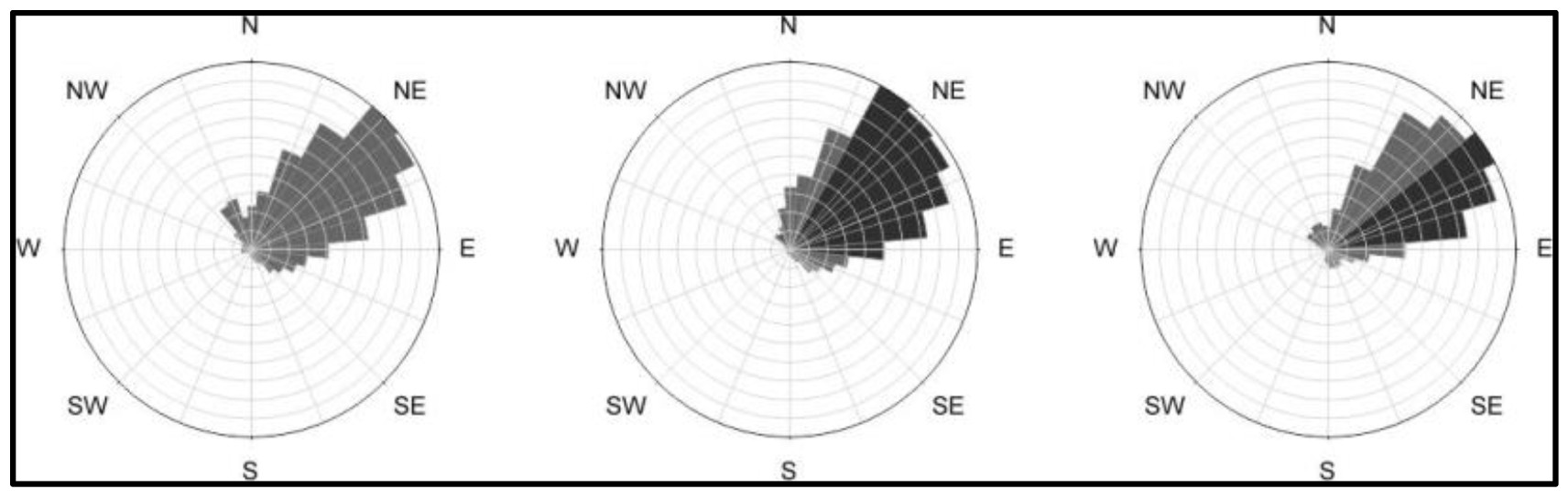
Typical wind fields in the Control plot. The wind roses shown are for 10:00-10:30 at the Control plot on August 29, 2014. Left to right is the CS, CD_1_, and CD_2_. Compass directions indicate the direction the wind was blowing *towards*. Solid black indicates a mean wind speed of 3 m/s.

Figure 10 shows examples of wind roses collected at the Forest plot from 13:30-14:00 on August 20, 2014. The top row shows (left to right) the FS, FW_1.8_, and FW_12.8_ (middle of the walkup tower). The bottom row shows (left to right) FW_23.8_, FD_1_, and FD_2_. Compared to the Control plot, the wind is generally lower speed and disorganized, although the anemometer at the top of the walkup tower, which is above the canopy, mostly shows large eddies rolling by along the NW-SE direction, plus a brief gust to the North. The wind in the source field was almost completely random in direction. The two sonic anemometers in the forest (W_1.8_ and W_12.8_) recorded the lowest wind speeds, which is due to the forest canopy creating turbulence that dissipates the flow’s kinetic energy.

**Figure 10.**
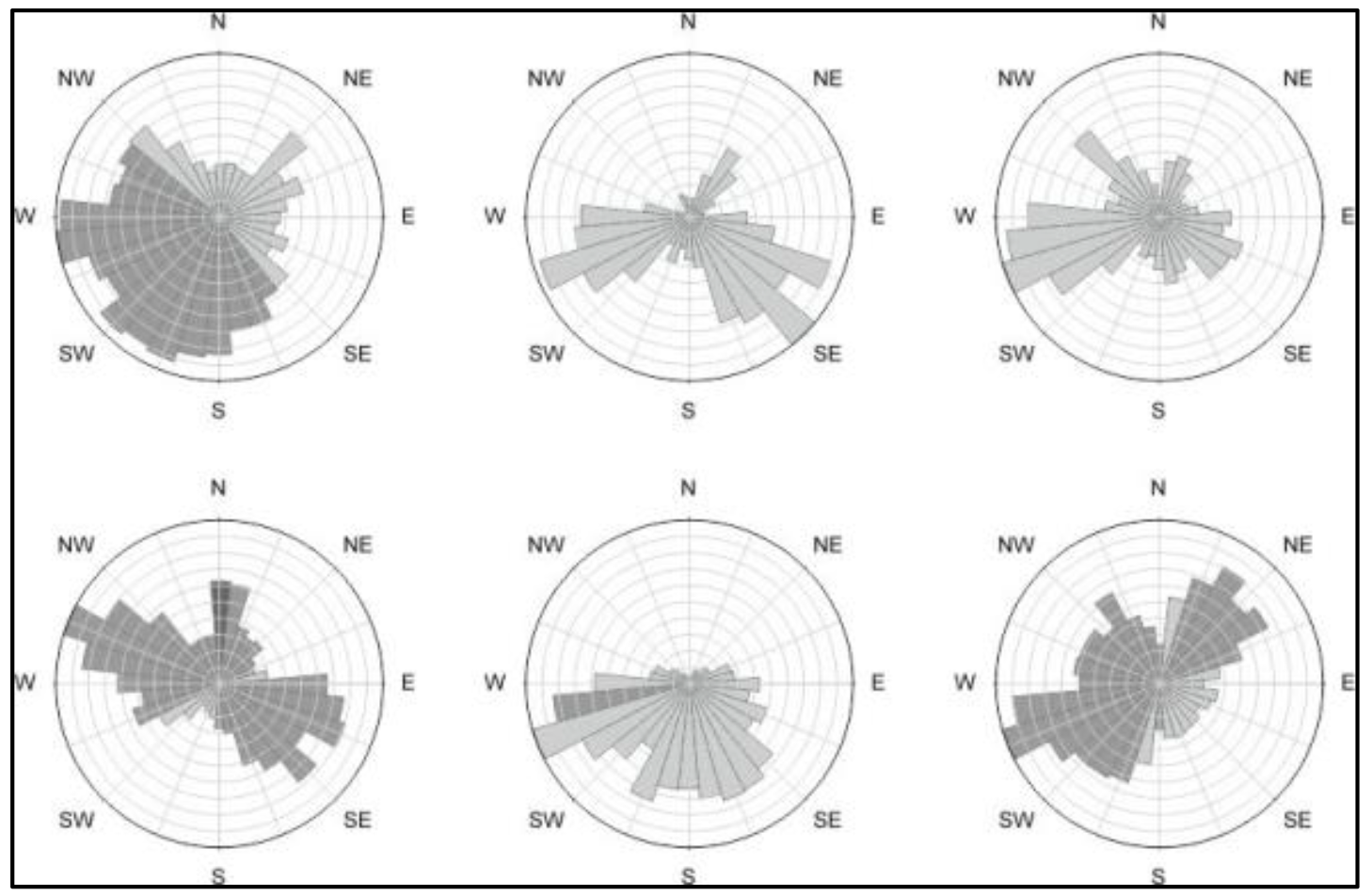
Typical wind fields in the Forest plot. Wind roses are shown for the Forest plot from 13:30-14:00 on August 20, 2014. The top row shows (left to right) the FS, FW_1.8_, and FW_12.8_ (middle of the walkup tower). The bottom row shows (left to right) FW_23.8_, FD_1_, and FD_2_. Compass directions indicate the direction the wind was blowing *towards*. Solid black indicates a mean wind speed of 3 m/s.

## 4. Discussion

Pollen source strength (PSS) is one of the most important factors in pollen-mediated gene flow, and our estimate for switchgrass PSS is the first in the literature. The theoretical maximum value for switchgrass PSS in our field plots was based on the number of pollen/anther and other quantified morphological traits, so the value 2.83 × 10^12^ must be considered an upper bound that would probably not be realized due to biotic factors (e.g. herbivory) and abiotic factors (e.g. wind conditions, humidity, or rain). In fact, pollen capture methods in this study showed that approximately 1/11^th^ of the theoretical maximum was advected above the switchgrass panicles. Previous PSS studies have mainly focused on maize pollen [Jia et al, 2007, Augspurger and Franson, 1987, Timerman et al, 2014, Sofiev and Bergmann, 2012, Arritt et al, 2007, Jarosz et al, 2005, Jarosz et al, 2003, Aylor, 2003]. Switchgrass (5500 pollen/anther) had about twice as many pollen/anther compared to maize (2000-2500 pollen/anther) [Sturtevant, 1881, Goss, 1968, Wallace and Bressman, 1949]. A study in Spain measured pollen/anther in a variety of grass species [Prieto-Baena et al, 2003]. The 5500 pollen/anther observed in switchgrass was higher than the reported values for 32 grass species in the study, but similar to two grasses (*Cynosurus echinatus*, 6745; *Elymus repens*, 5410), and less than four grass species (*Arrhenatherum album*, 12,045; *Festuca arundinacea*, 8941; *Lolium rigidum*, 10,453; *Sorghum halpense*, 7140). Although many factors are involved in pollen-mediated gene flow, the PSS value suggests that the likelihood of gene flow from switchgrass fields exceeds most other grasses studied.

Our central hypothesis about the ability of a narrow forest windbreak to mitigate downwind pollen dispersal was strongly supported by the case studies. However, the data do not provide direct information about the three-dimensional paths taken by the wind-blown pollen or how much pollen was captured by surfaces in the forest (e.g. leaves, stems, branches). Additional field work and the development of computational fluid dynamics models are needed to understand the advection and dispersal of pollen. Ultimately, it should be possible to optimize windbreak design (e.g. tree species, tree spacing) to create practical, low-cost barriers that will reduce pollen-mediated gene flow while promoting coexistence and ecosystem services.

## Acknowledgements

We are grateful to: Patrick Lienin (Univ. of Connecticut) for fieldwork and assistance with data processing; Junming Wang (Univ. of Illinois at Urbana-Champaign) for help with rotorod boxes; David Miller and Geoffrey Ecker (Univ. of Connecticut) for advice on many subjects; Jinwon Chung and Richard Rizzitello (Univ. of Connecticut) for assistance in data collection; and the farm crew for their work at the University of Connecticut Plant Science Research Farm. This research was supported by Biotechnology Risk Assessment Program Competitive Grant no. 2011-02189 from the USDA National Institute of Food and Agriculture to Carol Auer and Thomas Meyer. Support was also provided by the Storrs Agricultural Experiment Station and the University of Connecticut.

## Footnotes

1 Summation, not numerical integration, is correct here because the pollen was released during the daytime only, not continuously. In fact, numerical integration (trapezoidal method) produces a PSS estimate of 161 × 109, an overestimate of 20 × 109.

